# Characterization of diatom communities in restored sewage channels in the Boye catchment, Germany

**DOI:** 10.1101/2024.07.03.601863

**Authors:** Ntambwe Albert Serge Mayombo, Lisa Haverbeck, Mimoza Dani, Michael Kloster, Danijela Vidakovic, Bánk Beszteri, Andrea M. Burfeid-Castellanos

## Abstract

Restoration programs were initiated in different stream systems within the Boye catchment in the early 1990s to stop and reverse the negative impacts of anthropogenic disturbances. However, our knowledge of the effects of restoration works in this river network, specifically on benthic diatom communities, is still limited. Diatoms were used successfully to assess the impact of restoration works in a range of river networks around the world but received less attention in the Boye catchment. This study aimed to characterize benthic diatom communities in this restored river catchment, using digital microscopy methods. We collected samples in the spring and summer of 2020 at sites restored in different years (1995, 2005, 2010, 2012, and 2013) within the catchment. Our results showed no effect of season, restoration date (year), and location of the site along the streams on the most dominant diatom species. However, some rare taxa indicated significant variations between the seasons. Overall water quality in the streams ranged from moderate to very good, indicating the positive impacts of restoration works conducted in these former sewage channels.

## Introduction

Anthropogenic activities have exacerbated the degradation of river catchments around the world, reducing their capacity to deliver the most basic goods and services [1,2]. These essential ecosystems are frequently exposed to a rising number of stressors [3,4] and remain among the most substantially altered, resulting in unprecedented biodiversity loss [5,6]. River restoration initiatives generally seek to halt and then reverse the historical trends of freshwater biodiversity depletion caused by river channel deterioration, restoring them to their natural condition by enhancing their ecological health status [7–9]. Despite the fact that restoration is an improvement method, it is undeniable that it also introduces some level of disturbances in unclean water bodies [7,10,11]. However, our understanding of the effects of river restoration is still limited due to the fact that just a few projects have included comprehensive monitoring.

One of these projects include the rivers of the Boye catchment [12–14]. The Boye catchment is located in the densely populated and industrialized Ruhr Metropolitan Area, and they has been severely impacted by many anthropogenic stressors [12]. Previously, untreated wastewater from urban (e.g. open sewage channels), agricultural (e.g. surface runoff from farmlands), and industrial (e.g. coal mining) activities was discharged into these stream channels [12–14]. The morphology of the majority of these streams was altered since most of them were straightened, corseted in concrete and deepened. Additionally, previous intense coal mining activities in this region resulted in subsidence issues in large areas within the catchment [15].

Restoration programs were initiated in different stream systems within the Boye catchment in the early 1990s to stop and then reverse the negative impacts of anthropogenic disturbances [1], promote holistic management of these urban freshwater ecosystems, and comply with the European Framework Directive requirement of achieving good ecological health status [2,16], by the extended deadline of 2027. Therefore, an assessment of the success of recent restoration works within this watershed by evaluating the response of various biological quality components (e.g. fishes, macroinvertebrates, macrophytes, phytobenthos/phytoplankton; see [2] has become a necessity to determine whether all of the effort put in was worthwhile.

In stream ecosystems, diatoms are the most common, diverse, and best-studied group of biofilm-forming microalgae [17]. Their community composition reflects the state of their environment’s ecology, with quick changes responding to water composition variations, making them ideal candidates for biomonitoring in a variety of aquatic habitats [18]. Diatoms revealed successfully the impact of restoration activities in a range of other river systems throughout the world [11,19–23].

Some studies have investigated the response of biota, particularly macroinvertebrates, to the recent restoration efforts within the Boye watershed [12,14,24,25]. However, microphytobenthic communities, such as diatoms received less attention in these river systems. The goal of this study was to close this gap by characterizing diatom communities in restored streams within the Boye catchment, using virtual slides prepared as described in details in [26,27], using Biigle v.2.0 [28,29] This is the first study to present and discuss diatom community composition in these river systems as a function of the effects of time since restoration (year), sampling season (spring or summer), and site location (upstream, midstream, or downstream).

We hypothesize that we will be able to observe clear changes in diatom abundance and community compositions between sites within the Boye catchment based on sampling season, year of restoration, and location along the streams.

## Materials and methods

### Study site

For this investigation, we collected benthic diatom samples from various streams in the Boye catchment (Figure 1). These streams were once open sewage channels, but they have been restored over time, with some dating back to the early 1990s. Diatom biofilms were sampled in spring (April 24, 2020) and summer (July 2, 2020). Water temperature recorded *in situ* during sampling was 7.6 °C in spring (range from 6 °C to 10.7 °C) and 15.77 °C in summer (ranging from 15 °C to 17.8 °C) (Table 1). Conductivity ranged from 680 µS/cm to 1980 µS/cm, with a mean of 1021 µS/cm in spring, and 425 µS/cm to 1124 µS/cm, mean 805 µS/cm in summer (Table 1). Water pH ranged from 7.67 to 8.14 with a mean of 7.92, and from 7.26 to 7.98 with a mean of 7.65, in Spring and Summer respectively (Table 1). Only temperature and pH showed statistically significant differences (ANOVA *p* < 0.05) across seasons.

**Figure 1:**
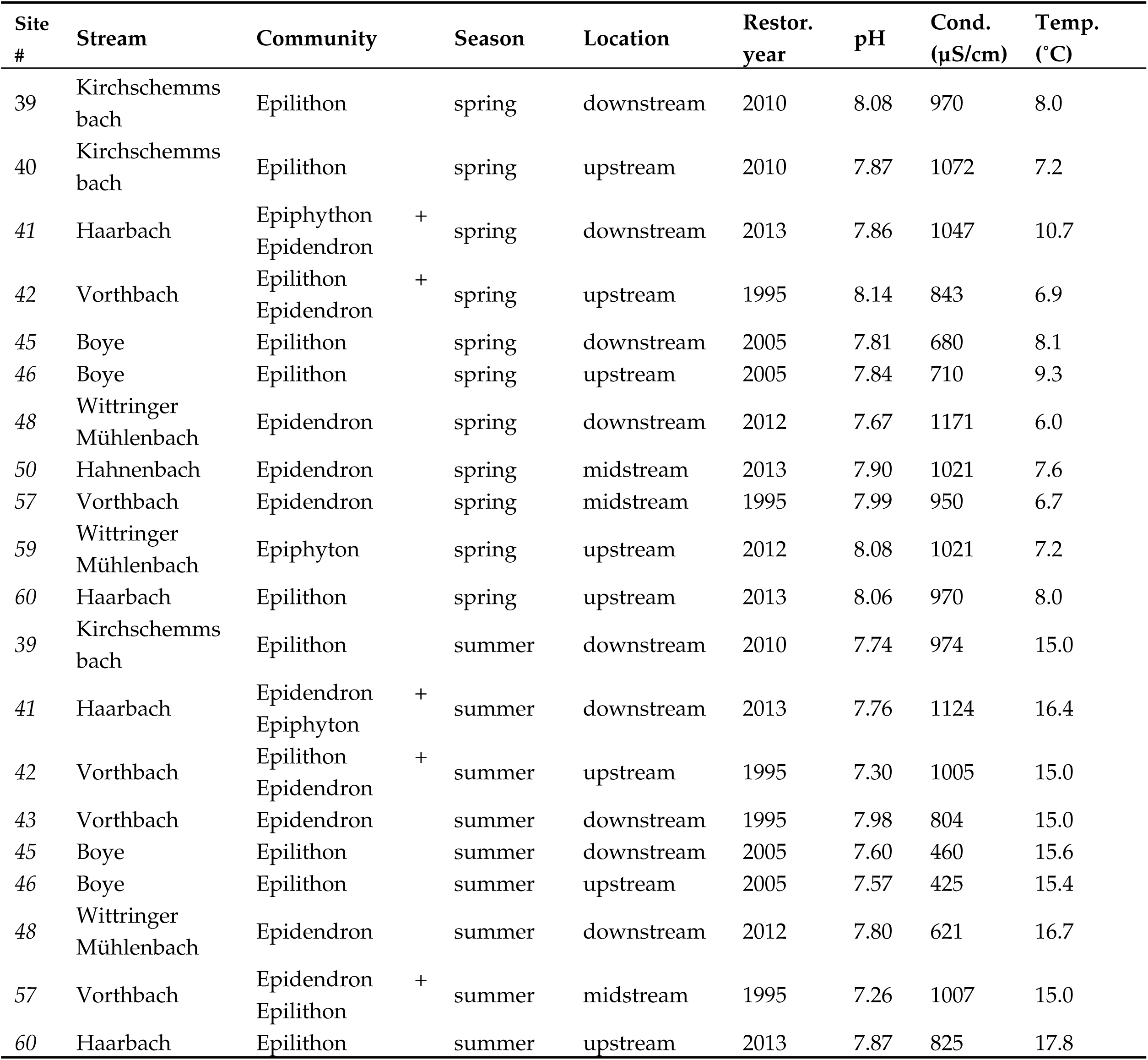
Map of the Boye catchment in Germany, the sampling sites are marked with green dots and numbered (more details in Table 1).

**Table 1.**
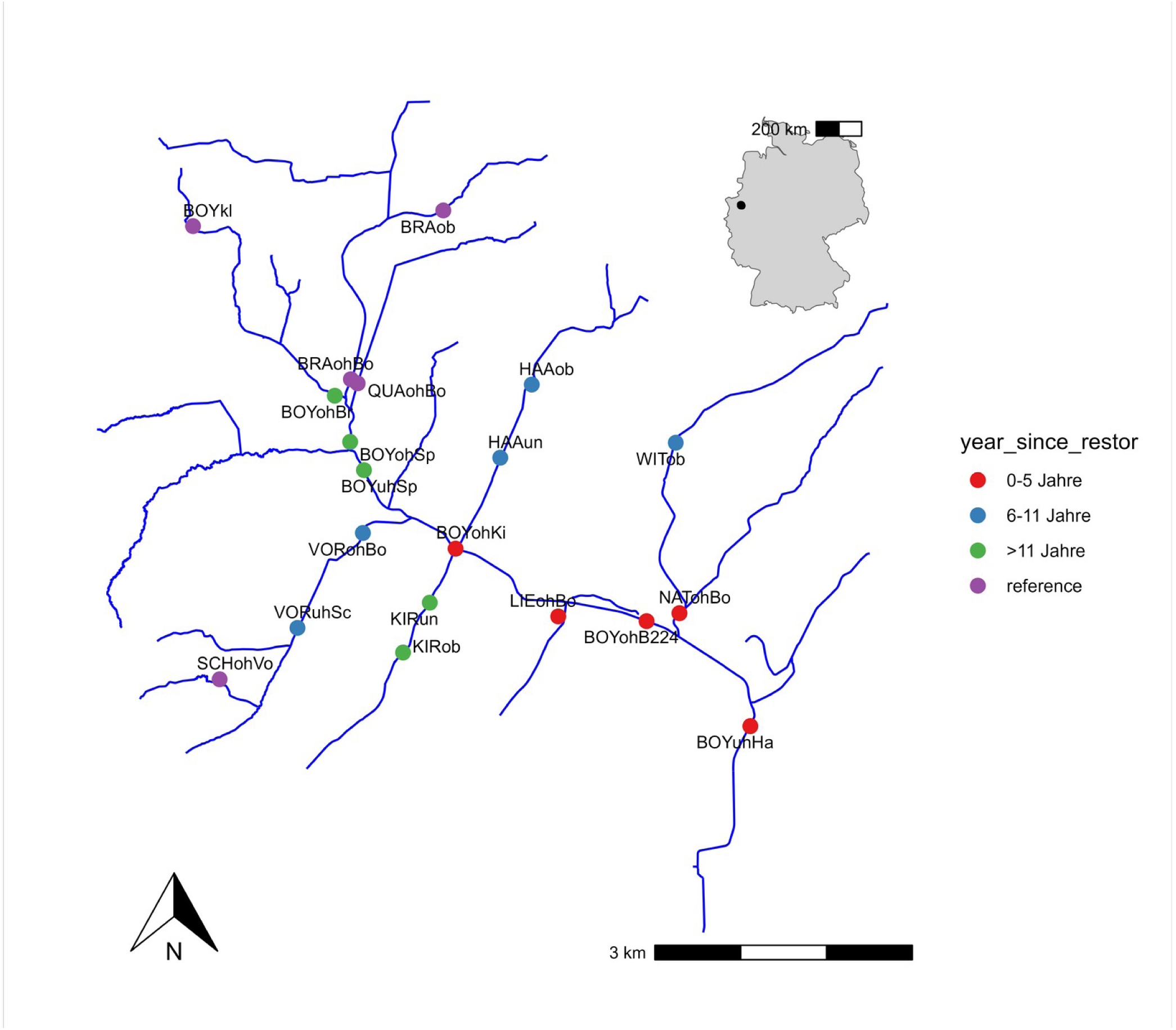
Sampling site details.

### Diatom biofilm sampling, processing and digital microscopy

Biofilm samples were collected by scrubbing a surface area of about 20 cm^2^ of at least three cobbles with a tooth brush. In sites where we were unable to find stones (epilithon), other hard substrata such as woods (epidendron) or macrophytes (epiphyton) were sampled. More details about the sampling sites are given in table 1. Immediately after collection, the samples were fixed in a solution of ethanol and then taken to the laboratory for further processing and analyses. For qualitative and quantitative analyses of diatom communities, we treated 10 mL of each sample with 30 ml of concentrated hydrogen peroxide (H_2_O_2_) on a hot plate, before adding few drops of concentrated hydrochloric acid (HCl) as described in Taylor et al. [30]. Then, a drop of the sample was mounted on a microscope permanent slide using Norland Optical Adhesive 61 (NOA 61). Finally, we scanned permanent slides with an Olympus Vs200 slide scanner (Olympus, Tokyo, Japan) at 63x magnification apochromatic and 1.24 Numeric Aperture to get virtual slides following the protocols detailed in Kloster et al. [26,27]. Diatoms on digital slide were identified by morphometric observations on Biigle 2.0 [28,29], using standard literature [31–33]. Diatom abundance data were exported as excel sheet for further analyses. We used the OMNIDIA software [34] for the calculation of diatom indices, such as the Specific Pollution Sensitivity Index (IPS) [35] and the Biological Diatom Index (IBD) [36] for water ecological health status assessment. The four-letter code devised for this software is used on diatom taxa to clarify multidimensional scaling.

Describe guilds and sense of them

### Statistical data analysis

We conducted all multivariate statistical analyses in the open-source statistical software R (v4.1.1) [37] using the ‘vegan’ package (v2.5-7) [38], and the ‘ggstatsplot’ package [39]. As proposed by Anderson et al. [41], diatom abundance data were log-transformed (log b (x) + 1 for x > 0, where b is the basis of the logarithm, here we used base 2) to minimize the weight of taxa with high abundance relative to those with low abundance to achieve multivariate normal distribution. All taxa with abundances less than 5% were put under a single label as “Others”. We ran the vegan ‘betadisper()’ function to calculate differences in multivariate homogeneity of group dispersion (variances or average distance to centroids) based on the distance matrix, then used the vegan ‘adonis()’ function to run a Permutational Multivatiate Analysis of Variance (PERMANOVA) to see how time since restoration, sampling season, and site location affected benthic diatom community composition. In order to visualize associations between samples based on distance matrices, non-metric multidimensional scaling (nMDS) was used with the vegan ‘metaMDS()’ function and the Bray–Curtis similarity index with 9999 permutations.

## Results

### Diatom community composition and diversity

In this study, we identified 232 distinct diatom taxa, from 52 genera. Only 31 taxa had abundance values greater than or equal to 5% (Figure 2, Table 2). The most dominant (abundance greater than or equal to 5 %) and frequent (occurring in at least 4 samples) diatom taxa in both sampling seasons were *Planothidium lanceolatum* (Brebisson ex Kützing) Lange-Bertalot (OMNIDIA code: PTLA), *Achnanthidium minutissimum* (Kützing) Czarnecki (ADMI), *Rhoicosphenia abbreviata* (C.Agardh) Lange-Bertalot (RABB), *Planothidium frequentissimum* (Lange-Bertalot) Lange-Bertalot (PLFR), *Cocconeis placentula* Ehrenberg (CPLA),*Gomphonema parvulum* var. *parvulum* (Kützing) Kützing (GPAR), and *Achnanthidium saprophilum* (Kobayasi et Mayama) Round & Bukhtiyarova (ADSA) (Figure 2). Each species is mostly found in a part of the biofilm.

**Figure 2:**
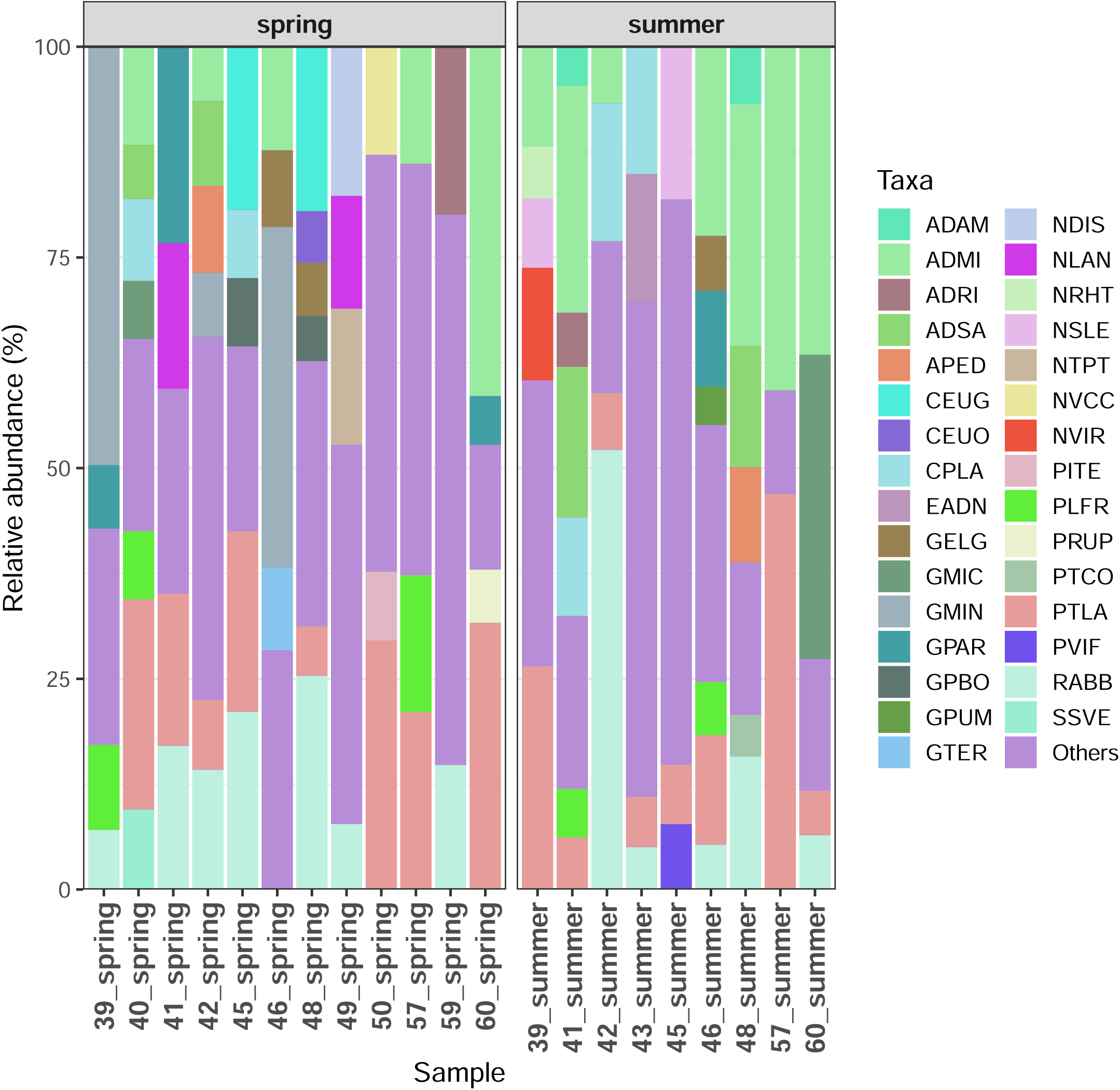
Relative abundance of dominant diatom species at different sites within the Boye catchment in spring and summer 2020.

**Table 2:**
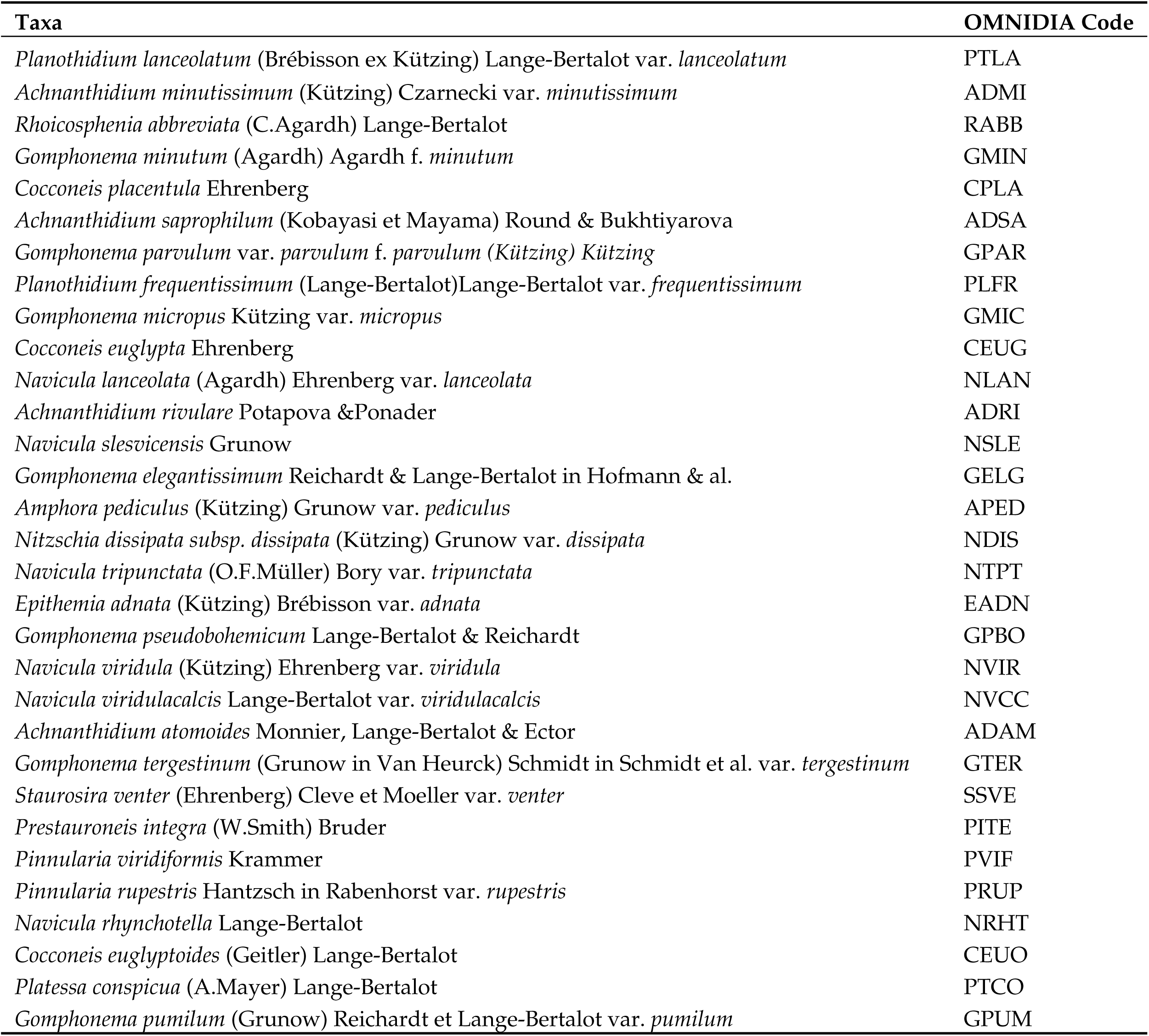
Most dominant diatom taxa identified in this study in decreasing order of abundances.

Despite the fact that the low profile guild predominated in high proportions in both seasons, we observed a significant rise in motile guild relative abundances in the spring, indicating that season had a major impact on the motile guild (ANOVA *p* < 0.001, Figures 3&4C). All other diatom guilds, however, did not show a significant effect of season (ANOVA *p* > 0.05, Figures 4A, B, D, E&F). The high profile guild, on the other hand, showed a significant influence of time since restoration (ANOVA *p* = 0.04).

**Figure 3:**
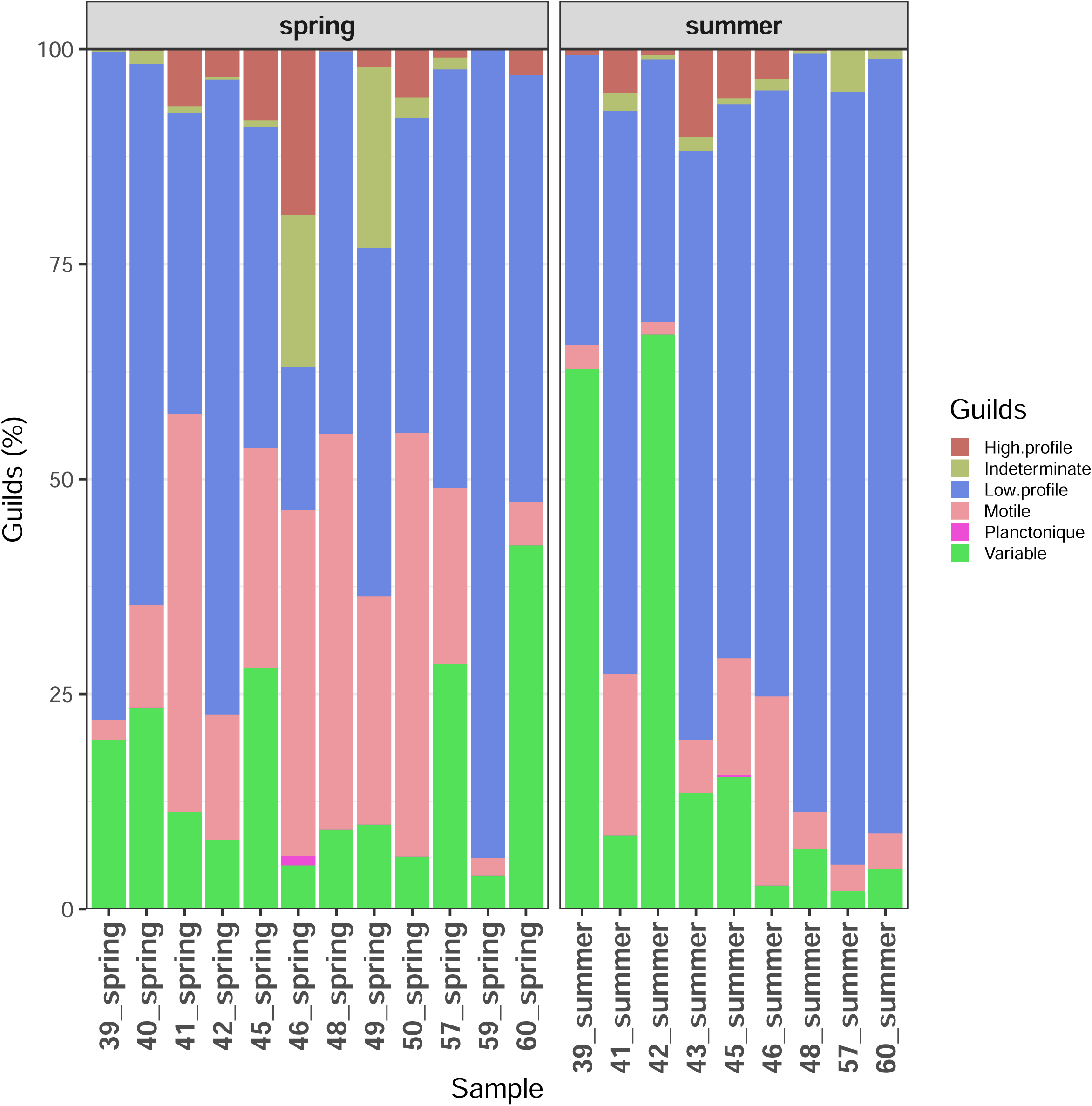
Relative abundance of different diatom guilds at different sites within the Bye catchment in spring and summer 2020.

**Figure 4:**
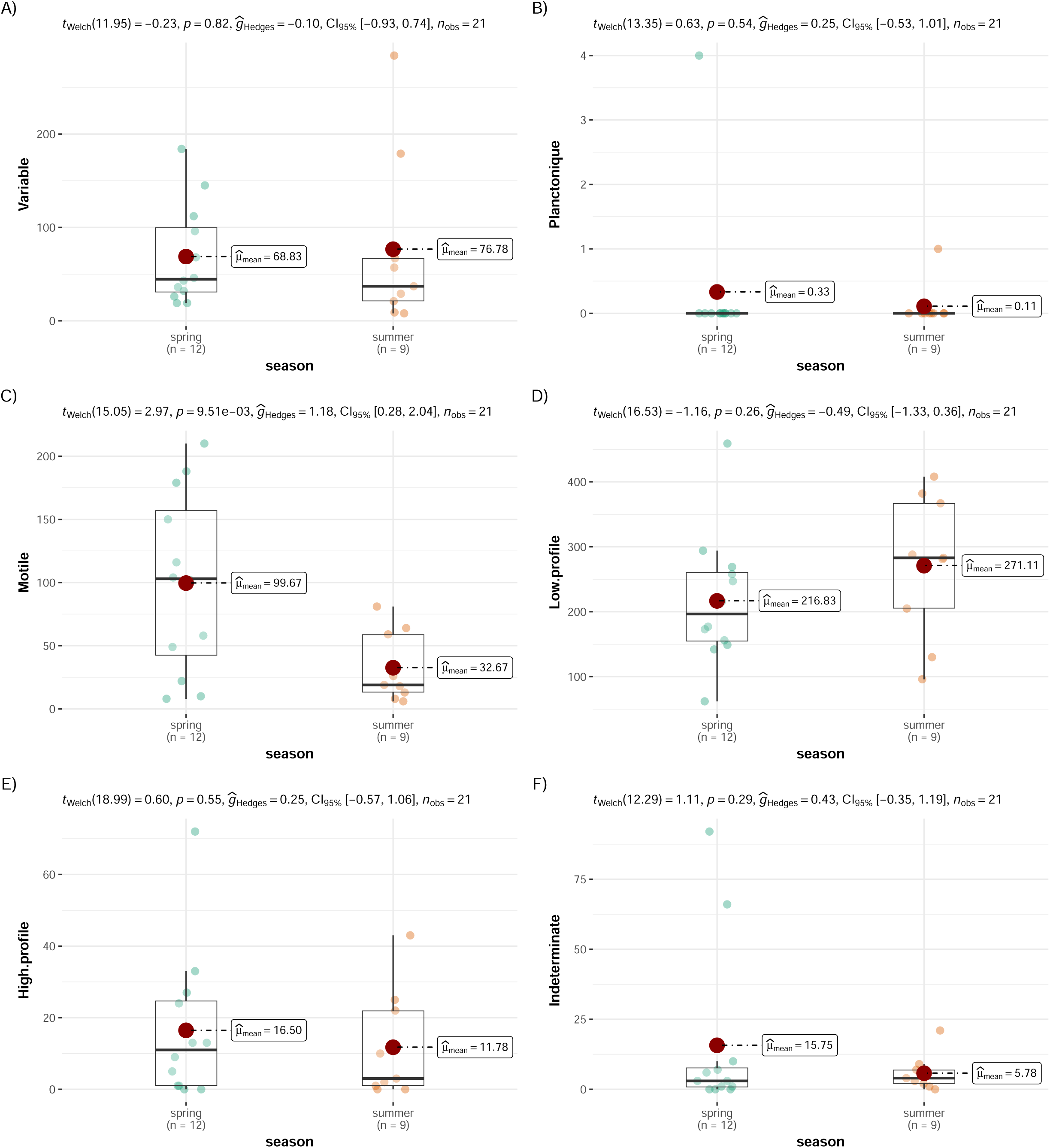
Variable (A), Planktonic (B), Motile guild (C), Low profile guild (D), High profile guild (E), and Intermediate (F) in spring and summer 2020. Results of ANOVA tests: p < only for motile guild, p > 0.05 for all the other guilds.

During the spring season, species richness (S) ranged from 25 taxa in site 39 (Table 1), to 71 taxa in site 50, (Figure 5A). In the summer, we observed low species richness (22 taxa) downstream Kirchschemmsbach and high species richness (61 taxa) downstream the Boye at sites 39 and 45, respectively (Figure 5A). Shannon-Wiener diversity index (H) ranged from 1.31 in site 59 to 3.55 in site 50 during the spring, with a mean of 2.54, and from 1.75 at site 48 to 3.26 at site 45 downstream the Boye in summer, with a mean of 2.49 (Figure 5B). Simpson diversity index peaked at 17.85 at site 50, and it was lowest 2.58 at site 59, upstream Wittringer Mühlenbach, in the spring (Figure 5C). In Summer, the greatest reading was 15.63 at site 45, and the lowest reading was 3.21 at site 48, downstream (Figure 5C). Pielou’s eveness (J) ranged from 0.40 (site 59 upstream Wittringer Mühlenbach) to 0.83 (site 50) and from 0.55 (site 48) to 0.80 (site 46), in spring and summer respectively (Figure 4D). However, we failed to observe any substantial difference between both seasons for all these diversity indices (ANOVA *p* > 0.05, Figures 5A, B, C&D). Additionally, none of the studied factors had a significant effect on the Specific Pollution Sensitivity (IPS) Index. The index showed values ranging from 12.8 to 18.3 (mean 15.13) in spring, and 13.7 to 16.2 (mean 14.99) in summer. The Biological Diatom Index (IBD), on the other hand, showed significant changes between seasons (Anova *p* = 0.04). Overall the index ranged 12.4 to 16.4 (mean 14.8) and 15.4 to 18.1 (mean 16) in spring and summer respectively.

**Figure 5:**
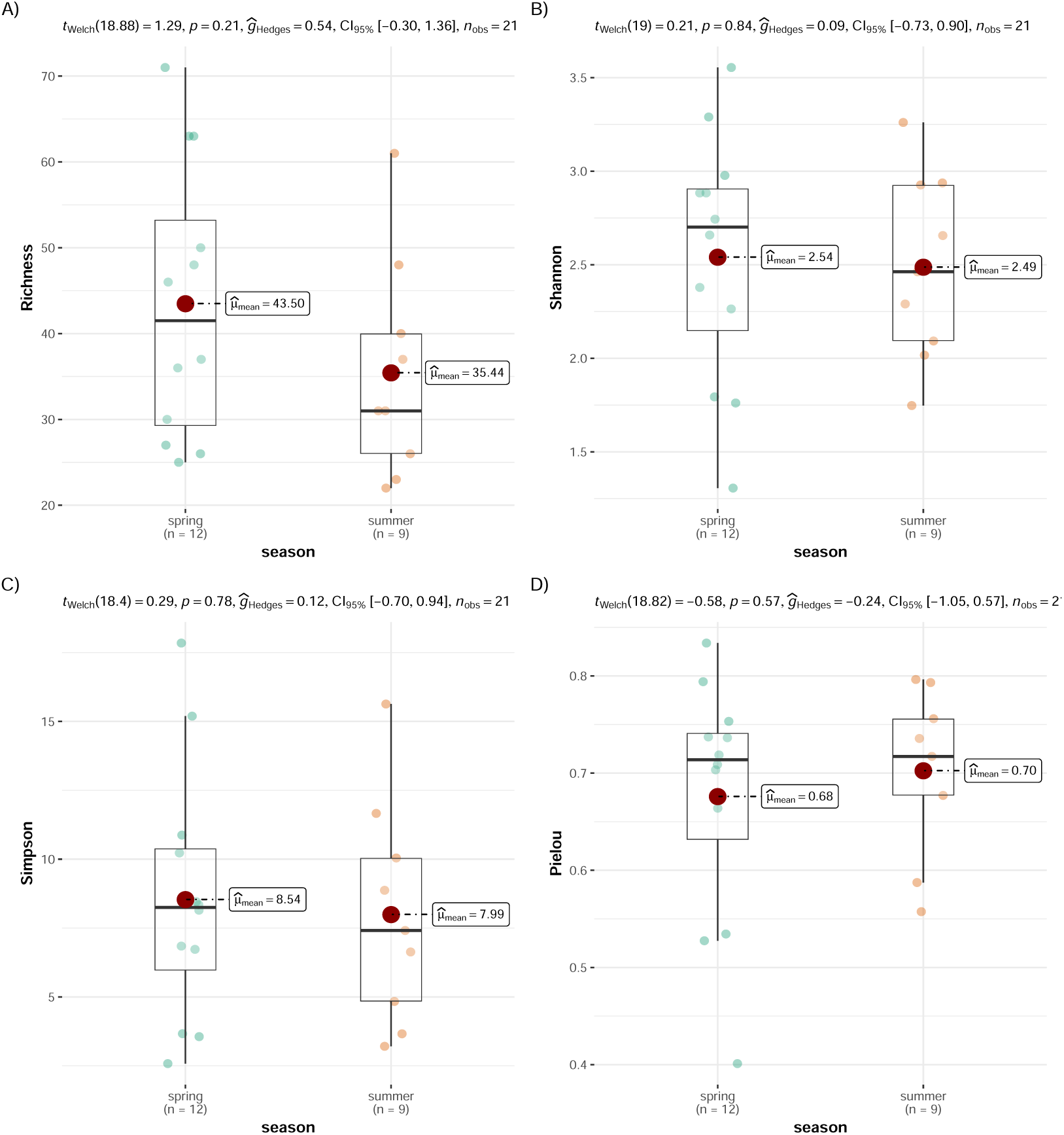
Species Richness (A), Shannon-Wiener diversity index (B), Simpson diversity index (C), and Pielou’s evenness (D) in spring and summer 2020. Results of ANOVA tests: p > 0.05 in all the cases.

### Effects of sampling season, time since restoration and site location on diatom communities

The test for multivariate homogeneity of group dispersion (distance from centroids) on our diatom species abundance dataset revealed that all groups were homogenous (*p* > 0.05). PERMANOVA run on a dataset which included only species with abundance values greater than or equal to 5% in each sample, did not demonstrate any significant differences between neither sampling season, duration of time after restoration, site location along the streams, nor their interactions (*p* > 0.05). However, when we ran PERMANOVA on the entire dataset, which also included all rare taxa with abundances less than 5%, we found that only sampling season had a significant effect on diatom abundances (Sum of Squares (*SS*) = 1.0658, *R^2^* = 0.229, *p* = 0.0017; Table 3). However, all two– and three-way interactions, as well as the effects of site location and time after restoration, were still not significant (*p* > 0.05, Table 3).

**Table 3:** Results of three-way crossed PERMANOVA of the effects of sampling season, site location, and time since restoration.

| Factor | df | SS | F | $R^2$ | $p^*$ |
| --- | --- | --- | --- | --- | --- |
| **Reduced dataset (only most abundant taxa)** |  |  |  |  |  |
| Season | 1 | 0.2460 | 0.9282 | 0.0529 | 0.5147 |
| Restoration_date | 4 | 0.8603 | 0.8117 | 0.1849 | 0.7064 |
| Site | 2 | 0.4674 | 0.8820 | 0.1005 | 0.5767 |
| Season x Restoration_date | 4 | 1.1162 | 1.0531 | 0.2399 | 0.4050 |
| Season x Site | 2 | 0.4908 | 0.9261 | 0.1055 | 0.5367 |
| Restoration_date x Site | 5 | 1.0232 | 0.7723 | 0.2199 | 0.7788 |
| Season x Restoration_date x Site | 1 | 0.1840 | 0.6945 | 0.0396 | 0.7592 |
| Residual | 1 | 0.2650 |  | 0.0570 |  |
| Total | 20 | 4.6528 |  | 1.0000 |  |
| **Whole dataset (with rare taxa)** |  |  |  |  |  |
| Season | 1 | 1.0658 | 6.0910 | 0.2291 | **0.0017** |
| Restoration_date | 4 | 1.1536 | 1.6482 | 0.2479 | 0.1545 |
| Site | 2 | 0.4357 | 1.2449 | 0.0936 | 0.2952 |
| Season x Restoration_date | 4 | 0.6925 | 0.9894 | 0.1488 | 0.4678 |
| Season x Site | 2 | 0.2956 | 0.8446 | 0.0635 | 0.6337 |
| Restoration_date x Site | 5 | 0.7342 | 0.8392 | 0.1578 | 0.6726 |
| Season x Restoration_date x Site | 1 | 0.1005 | 0.5743 | 0.0216 | 0.8827 |
| Residuals | 1 | 0.1750 |  | 0.0376 |  |
| Total | 20 | 4.6528 |  | 1.0000 |  |

\* The only significant ( $p < 0.05$ ) main effect of the season is highlighted in bold. df: degrees of freedom; SS: sum of squares; F: F-statistic, $R^2$ : variance explained by the groups, $p$ : p-values.

Indicator species analysis revealed that only *Achnanthidium minutissimum* (Kützing) Czarnecki var. *minutissimum* (ADMI) was the only most common taxa to have near significant (*p* = 0.045) differences in abundances between seasons, occurring in slightly higher numbers in summer. *Navicula cari* Ehrenberg (*p* = 0.02) and *Diploneis calcilacustris* Lange-Bertalot et A. Fuhrmann (*p* = 0.02) increased also significantly in abundances in summer, but they were classed as rare taxa together with all those that did not reach 5% in abundance. In the spring, *Hippodonta capitata* (Ehr.) Lange-Bertalot, Metzeltin and Witkowski (HCAP) and *Ulnaria ulna* (Nitzsch) Compere var. *ulna* showed nearly significant rise in abundance (*p* = 0,049 and 0.049, respectively). The nMDS plots revealed large seasonal overlapping and widely dispersed sample sets (Figures 6 A&B). Although the analysis yielded a high stress value (0.19), we observed a severe distortion in the arrangement of sample points scattered all over the two-dimensional (2D) space displayed on the graphs, despite the high correlation between the observed dissimilarity and the ordination distances (non-metric fit *R^2^* = 0.99, linear fit *R^2^* = 0.98, Figures 6 A&B).

**Figure 6:**
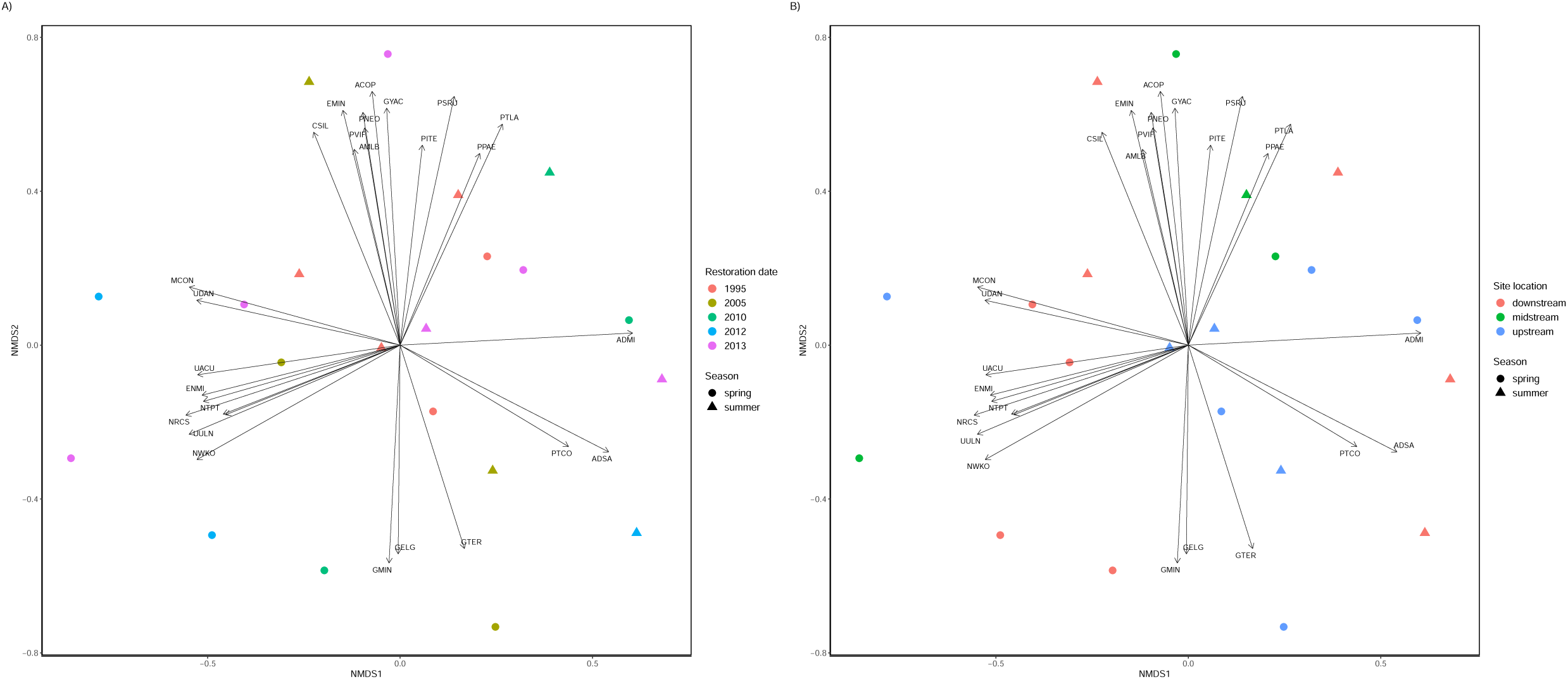
Non-metric multidimensional scaling (nMDS) graphs showing distances between diatom communities in samples based on (A) Restoration date and season and (B) site location and season.

## Discussion

In the current study, we expected to observe clear differences in diatom abundance and community compositions between sites within the Boye catchment based on sampling season, date of restoration (year), and location along the streams. Our data demonstrated that the dominant benthic diatom taxa identified in this watershed, except *Achnanthidium minutissimum* were unaffected by any of the investigated factors. The sampling season, on the other hand, had a significant effect on the full dataset, which included rare taxa. *A. minutissimum* was the only most dominant and frequent taxon to show nearly significant differences in abundances between the seasons. This is a pioneer diatom species [21,24], that occurs in large numbers at a variety of trophic levels [32]. It thrives abundantly in alkaline and acidic habitats, as well as in oligotrophic and hypertrophic waters [42]. Several studies reported *A. minutissimum* to be [11,19,21,43–45] an indicator of good water quality, for example low nutrients content.

This diatom species was the most numerous and important, contributing to differences between sites based on restoration status in “boreal rivers” [21]. We speculate that the increase in abundance of *A. minutissimum* in Summer 2020 was an indication of better in water quality during this season. *Navicula cari* and *Diploneis calcilacustris*, the other two species that showed substantial seasonal differences, increased in relative abundance in summer, albeit they remained scarce.

All the other most prevalent diatom species were found in roughly the same amounts in both seasons, with no notable variations between them. Most of these taxa, such as *Planothidium lanceolatum*, *Rhoicosphenia abbreviata*, *Gomphonema minutum*, *Cocconeis placentula*, *Gomphonema parvulum* var. *parvulum*, *Planothidium frequentissimum*, *Gomphonema micropus* and *Achnanthidium saprophilum* have a wide range of tolerance to various ecological conditions. For example, *P. lanceolatum* and *P. frequentissimum* very often co-occur in electrolyte-poor to rich waters, as well as in oligotrophic to polytrophic, up to β-mesosaprobic waters [31,32]. *Rhoicospehenia abbreviata* and *Cocconeis placentula* were reported as characteristic indicator species of sites with intermediate level of pollution [46]. These dominant diatom species were also found in restored reaches with better water quality in the Bzura River, Poland [19]. The presence of these taxa in high abundance suggests some improvements in water quality within the Boye catchment even though there are still some levels of various anthropogenic stressors that could be entering the system as the waters flow through the densely populated urban areas.

Our dataset showed a clear dominance of low profile diatom ecological guild in both seasons. Low profile guild increase in relative abundance in nutrients-poor and high disturbances (e.g. high flow velocity) conditions [47,48]. Our results revealed no effect of neither season, restoration date, nor site location on low profile guilds. Trábert et al. [49] found no correlation between low profile guild abundance and any of the investigated factors, such as nutrients and environmental variables, in the Danube. The relative abundance of motile guild increased significantly throughout the spring season. Motile guild diatom species are more nutrients-dependent and susceptible to perturbations [47,48]. We speculate that water quality in the streams degraded slightly during the spring season, which favoured motile guild diatoms to thrive. This degradation is also seen through the slight increase in conductivity and pH observed during this season.

The Biological Diatom Index (IBD) [36] revealed significant differences between the spring and summer seasons, indicating moderate to good ecological status in spring, and even higher ecological status in summer. Therefore, in the spring, water quality ranges from moderate to good, and in the summer, it rises to very good. The Specific Pollution Sensitivity Index (IPS) [35] on the other hand, was not responsive to any of the examined conditions but did indicate moderate to good ecological status in the spring and even higher ecological status in the summer. Similarly, according to IPS, water quality was satisfactory to good in the spring and improved to very good in the summer. These findings point to an increase in water quality within the Boye catchment, which has been repaired for at least seven years at some sites and even over a decade at others. According to PERMANOVA run on the most dominant taxa dataset, diatom community composition and abundances were not influenced by season, time since restoration and location of the site along the streams. However, season had a significant effect on the whole abundance dataset, including rare taxa. Our results imply that diatom community composition in these sites that we sampled were more or less similar, with no significant differences among them. We speculate that diatom communities in these sites evolved to much stable or climax communities over the years.

In conclusion, our results revealed no significant effects of season, time since restoration and location of the sites along the streams. The dominant diatom species in all the sites were very similar during spring and summer. Given the fact that all the sites we sampled in this study were restored for over at least seven years, it would be necessary? to also sample in sites that were restored more recently. This study is a baseline report of the ecological status of the rivers within the Boye watershed based on benthic diatom communities. Although we lack the knowledge of the conditions of these streams prior to the start of restoration works to assess their effectiveness at this point, these baseline data will be helpful for future monitoring. These results will be important for the ongoing research under the Collaborative Research Centre (CRC) 1439 RESIST, in which we are investigating the functional and compositional responses of stream microphytobenthic communities to multiple stressors increase and decrease within the Boye catchment using both field work and large scale ExStream mesocosm experiment.

